# Allograft Inflammatory Factor-1 regulates immune activation states and is required for effective pathogen-specific T cell immunity during infection

**DOI:** 10.64898/2026.07.23.739600

**Authors:** Lais Rekowsky, Ricardo Louzada da Silva, Ayane Resende, Jonathan Seenarine, Marissa Macchietto, Tatiana de Moura, Diana M. Elizondo, Michael W. Lipscomb

## Abstract

Allograft inflammatory factor-1 (AIF1) is a scaffold protein expressed predominantly in myeloid antigen-presenting cells (APCs) and associated with inflammatory disease. Although genetic studies have linked AIF1 loci to immune traits, its causal role in physiological immune responses remains poorly defined. We examined AIF1 deficiency using a conditional knockout model with deletion of AIF1 in hematopoietic cells (AIF1-cKO) during development and challenged mice with Listeria monocytogenes. AIF1 loss impaired bacterial clearance and diminished inflammatory responses, indicating reduced immune readiness during infection. AIF1-cKO mice exhibited impaired expansion of antigen-specific CD4+ and CD8+ T cells, accompanied by regulatory-associated phenotypic changes. These defects were associated with reduced cDC1 frequencies and transcriptional and phenotypic remodeling of splenic macrophages toward a less inflammatory state. Single-cell RNA sequencing revealed transcriptional alterations across multiple myeloid and lymphoid compartments despite AIF1 expression being largely restricted to myeloid APC, indicating broader immune remodeling. Increased Tgfbr1 expression was a recurrent feature across several immune populations. Consistent with this finding, AIF1-deficient cells displayed enhanced TGFβ responsiveness, while Tgfbr1 silencing partially restored inflammatory responses and T cell priming *ex vivo*. These findings establish AIF1 as a regulator of immune competence that promotes effective innate and adaptive immune responses.

## Introduction

Allograft inflammatory factor-1 (AIF1) is a conserved MHC class III-encoded calcium-binding scaffold protein^1-3^, expressed predominantly in myeloid antigen-presenting cells (APCs), such as macrophages and conventional dendritic cells (cDC)^4-9^. At the cellular level, AIF1 has been linked to cytoskeletal remodeling, phagocytosis and inflammatory signaling programs^10-13^, suggesting a role in modulating innate immune activation, inflammatory responses and T cell priming capacity^14-18^.

Consistent with these cellular functions, AIF1 has emerged as a recurrent candidate regulator of immune function through both human genetic studies and experimental investigations of inflammatory disease^8, 19-29^. Multiple studies have associated AIF1-linked loci or altered AIF1 expression with autoimmune, inflammatory and metabolic disorders, suggesting a strong role in regulating immune homeostasis and disease susceptibility^20, 21, 30-35^. Recent human translational evidence has demonstrated that a natural low-expressing AIF1 promoter variant restricts macrophage-driven tissue infiltration and protects against inflammatory pathology, establishing a direct link between genetic expression strength and myeloid APC activation capacity^36^. Additionally, experimental murine studies have implicated AIF1 as an important modulator of innate inflammatory responses and adaptive immunity^16, 17^.

However, the extent to which AIF1 contributes to the coordinated regulation of immune responses *in vivo* remains poorly defined. To date, studies have largely been limited to correlative human observations, reductionist *ex vivo* systems, or non-antigen-specific inflammatory models, leaving unresolved whether AIF1 is required for effective immune responses during infection. In particular, it remains unknown whether loss of AIF1 compromises the coordinated innate and antigen-specific adaptive immune responses required for effective pathogen clearance *in vivo*.

Here, we address this question using hematopoietic AIF1-deficient mice challenged with *Listeria monocytogenes* (LM) to examine immune responses during infection. We find that loss of AIF1 impairs bacterial clearance, reduces antigen-specific CD4+ and CD8+ T cell responses, and alters the myeloid APC composition and functional states. These defects are accompanied by transcriptional and phenotypic remodeling across myeloid and lymphoid compartments, indicating that loss of AIF1 in antigen-presenting cells is associated with broader remodeling of the immune environment.

Together, these findings identify AIF1 as a regulator of immune competence and demonstrate that AIF1 is required for innate inflammatory responses and effective adaptive T cell priming during infections. These studies provide direct *in vivo* evidence that AIF1 is an important determinant of immune competence and establish a functional framework for understanding how AIF1-associated genetic variation and dysregulation may influence immune-mediated disease.

## Results

### Generation of mice deficient for AIF1 in hematopoietic populations

To directly evaluate the role of AIF1 in hematopoietic immune cell function *in vivo*, a conditional AIF1 knockout mouse model (AIF1^fl/fl^) was generated and crossed to the Vav1iCre strain to produce hematopoietic-specific deletion of AIF1. Experimental cohorts consisted of Vav1iCre^+/-^AIF1^fl/fl^ mice (AIF1-cKO) and littermate Vav1iCre^+/-^AIF1^fl/+^ and AIF1^fl/fl^ internal controls. Importantly, no significant differences in baseline immune phenotypes were observed between Vav1iCre^+/+^AIF1^fl/+^ and AIF1^fl/fl^ littermate controls, supporting the use of both strains as appropriate controls (**Supplemental Figure 1A**).

Successful recombination of the floxed AIF1 allele and presence of the Vav1iCre transgene were confirmed by PCR genotyping across all experimental cohorts (**Supplemental Figure 1A**). Efficient deletion of AIF1 protein was confirmed across hematopoietic compartments, including total splenocytes, bone marrow-derived cells, macrophages, and cDC. AIF1 protein was readily detected in control animals but absent in corresponding AIF1-cKO populations (**Supplemental Figure 1B-C**), confirming effective hematopoietic ablation of AIF1.

### Pathogen challenge in AIF1-deficient mice results in impaired bacterial clearance and altered innate immune activation states

To determine whether AIF1 is required for effective host defense, hematopoietic AIF1 conditional knockout (AIF1-cKO) mice were infected with *LM*. AIF1-cKO mice exhibited significantly impaired bacterial clearance, with higher splenic pathogen burden over a 5-day infection course compared with control animals (**Figure 1A-B**). To assess transcriptional changes in the splenic immune compartment, key inflammatory and immunoregulatory genes were evaluated in total splenocytes. AIF1-cKO mice exhibited significantly increased Arg1 expression without a corresponding increase in Nos2, suggesting altered macrophage activation states during infection (**Figure 1C**). Flow cytometric analysis further demonstrated phenotypic remodeling of the splenic macrophage compartment in AIF1-cKO mice. The F4/80^+^CD11b^lo^ macrophage compartment showed reduced MHC class II expression together with increased expression of regulatory markers Mertk and CD206 (**Figure 1D**), marking a shift toward a less inflammatory and more immunoregulatory activation state. In parallel, DC composition was altered, with reduced frequency of CD24^+^CD172a^neg^ cDC1 subsets within the CD11c^+^MHC class II^+^ population (**Figure 1E**), as demonstrated prior under *ex vivo* conditions^17^.

**Figure 1.**
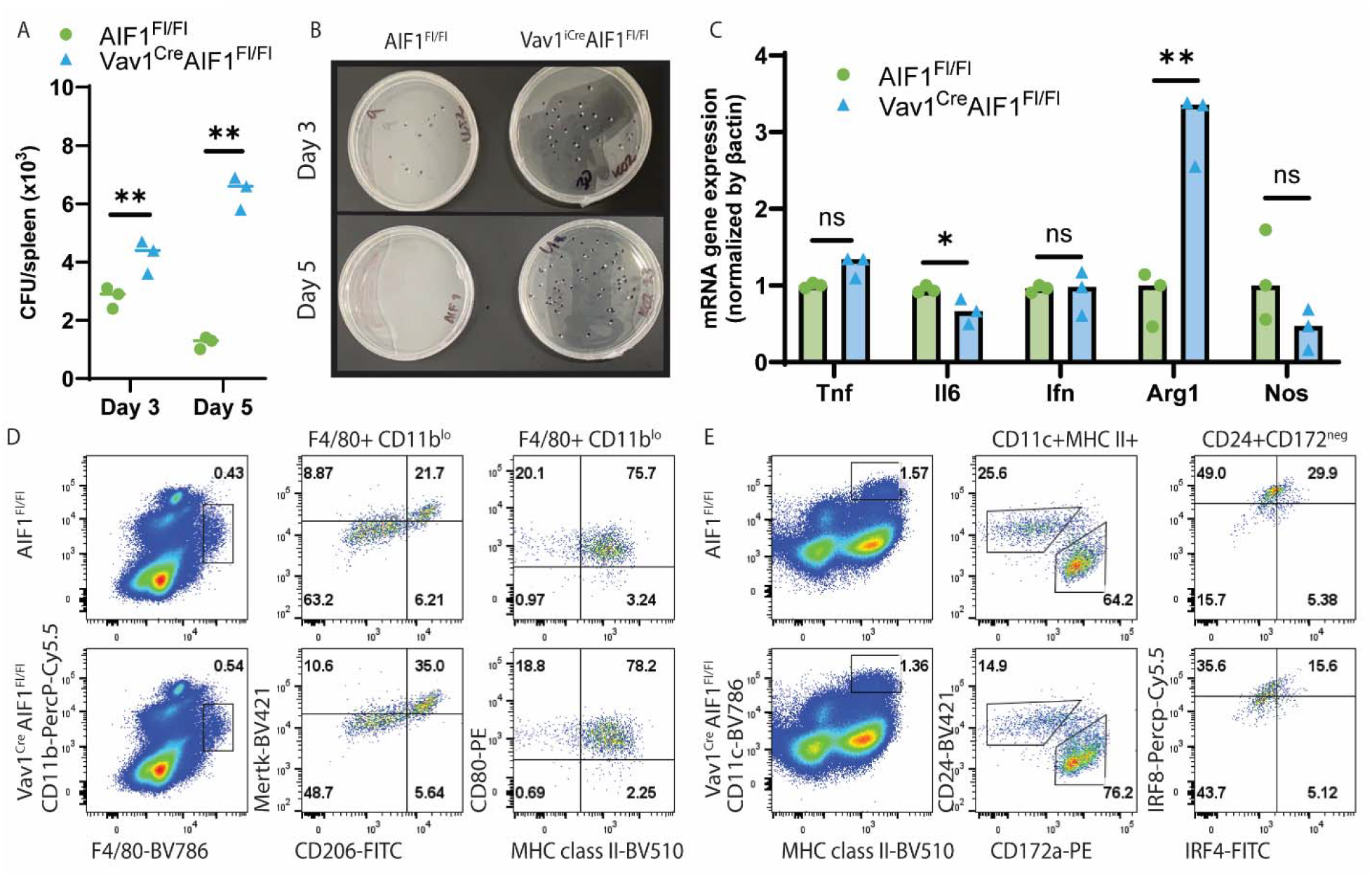
Pathogen challenge in AIF1-deficient mice results in depressed innate immune responses and clearance of microorganisms. (**A-B**) Splenic bacterial burdens (CFU) in WT (AIF1^fl/fl^) and AIF1-cKO (Vav1^iCre^AIF1^fl/fl^) mice at days 3 and 5 post intravenous LM-OVA infection (1×10^3^ CFU). (**C**) Isolated total splenocytes assessed by qPCR analysis for Tnf, Il6, Ifng, Arg1 and Nos2 mRNA levels from LM-infected mice at day 3. (**D**) Flow cytometric analysis of splenic F4/80+CD11b^lo^ macrophages showing frequency of Mertk+CD206+ and CD80^+^MHC class II^+^ in AIF1-cKO vs control mice post-infection. (**E**) Frequency of CD24+CD172a^neg^ cDC1 and CD24^neg^CD172a+ cDC2 subsets among splenic CD11c+MHC class II+ DC. Evaluation of IRF8 vs. IRF4 to confirm cDC1 signatures. Data are mean ± SEM (n=3-8 mice/group); statistical significance was determined by unpaired two-tailed Student’s t-test (*p<0.05, **p<0.01).

### AIF1 deficiency impairs *in vivo* antigen-specific T cell expansion and promotes regulatory CD8+ T cell programming during LM infection

To determine whether defective bacterial clearance in AIF1-cKO mice reflected impaired adaptive immunity, antigen-specific T cell responses were quantified following infection with *Listeria monocytogenes* expressing ovalbumin (LM-OVA). Four days after infection, the frequency of OVA-specific CD8^+^ T cells in the spleen, detected using H-2Kb_-SIINFEKL_ tetramers, was significantly reduced in AIF1-cKO mice compared with controls (**Figure 2A**). MHC class I-restricted tetramer specificity was confirmed by the absence of staining in CD4^+^ T cells. Beyond the reduction in antigen-specific CD8^+^ T cell expansion, the responding population exhibited a distinct regulatory-associated phenotype, characterized by increased co-expression of CD122 (IL-2Rβ) and PD-1 (Figure 2B), markers associated with regulatory or Tr1-like CD8^+^ T cell states.

**Figure 2.**
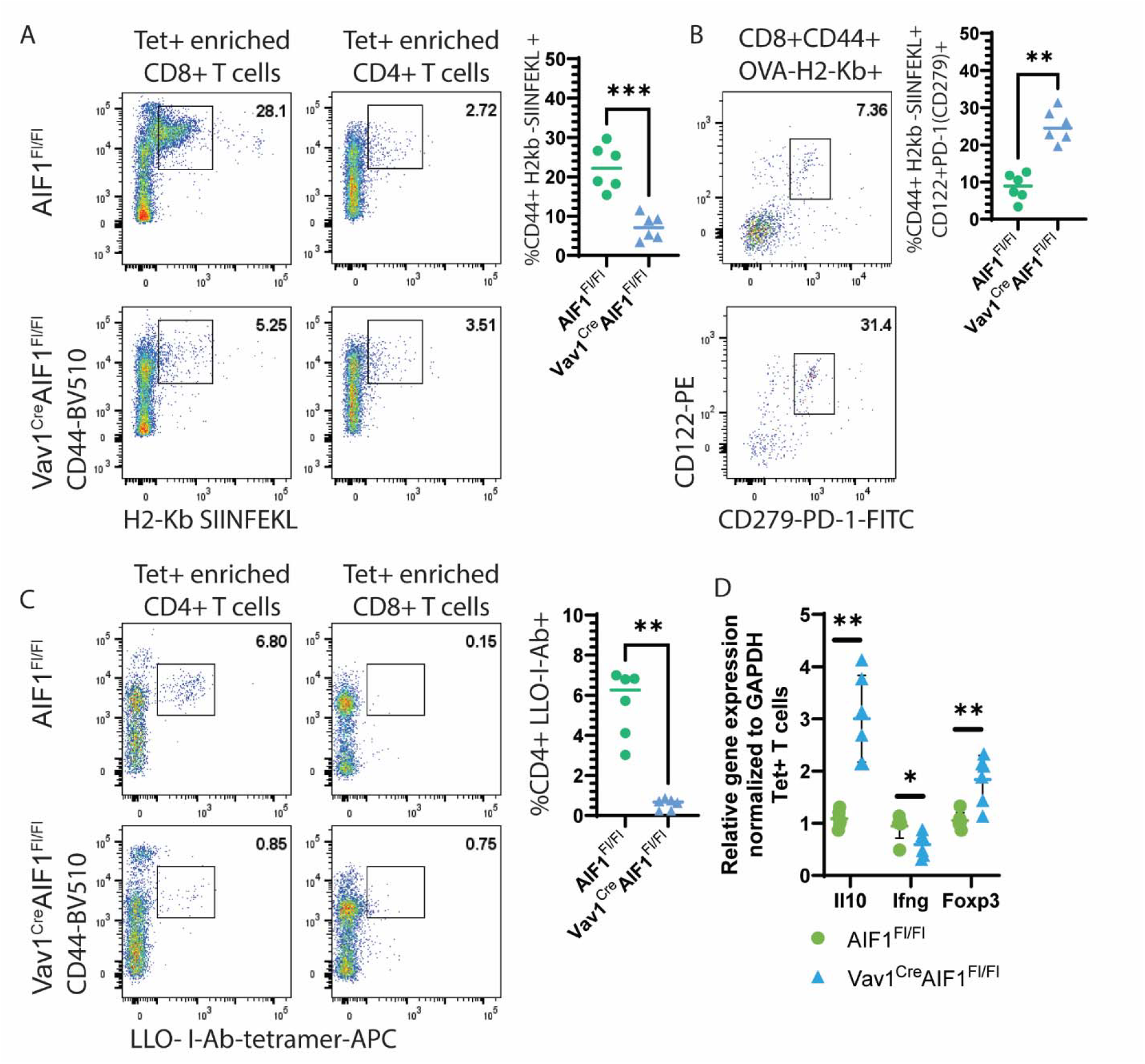
Loss of AIF1 restrains LM-specific CD8+ and CD4+ T cell expansion *in vivo* and shifts programming towards a regulatory state. (**A**) Frequency of OVA-specific CD8+ T cells (H-2KLJ-_SIINFEKL_ pMHC tetramer+) in spleen at day 4 post-LM-OVA infection; CD4+ T cells evaluated as internal non-MHC class-I restricted negative control. Isolated cells are enriched for tetramer+ cells using magnetic beads prior to flow cytometric analyses. (**B**) Frequency of CD122+PD-1+ cells among OVA-specific CD44+CD8+ T cells. (**C**) Frequency of LLO-specific CD4+ T cells (I-A□ pMHC tetramer^+^) in spleen at day 4 post-infection. Data are shown as representative dot plots with corresponding quantification of biological replicates (mean ± SEM, n=3-6 mice/group); statistical significance was determined by unpaired two-tailed Student’s t-test (*p<0.05, **p<0.01).

T helper cell responses were similarly affected, with a marked reduction in LM-specific CD4^+^ T cells in AIF1-cKO mice relative to controls, as measured using I-A^b^ tetramers recognizing the Listeriolysin O (LLO) epitope (**Figure 2C**). No tetramer binding was detected in CD8^+^ T cells, confirming MHC class II-restricted staining specificity. Because LM-specific CD4^+^ T cells were present at such low frequency in AIF1-cKO mice, antigen-specific populations were enriched by MACS-based tetramer-positive isolation prior to qPCR analysis. This analysis revealed increased expression of Foxp3 and Il10 together with reduced expression of Ifng, demonstrating a transcriptional shift away from inflammatory effector programs and toward regulatory-associated features (**Figure 2D**), paralleling prior *ex vivo* findings^16, 37^.

### Single-cell RNA sequencing analysis of AIF1-deficient leukocytes reveals altered transcriptional signatures in antigen-presenting cells and T cells

To define immune states associated with AIF1 deficiency, scRNA-seq was performed on spleens from AIF1-cKO and control mice at steady state. Integration and comparative analysis revealed no major differences in overall cell distribution (**Supplemental Figure 2A**). As expected, AIF1 expression was largely restricted to myeloid APC (**Supplemental Figure 2B-C**)^17^. Vav1 expression was detected across the hematopoietic populations analyzed, consistent with the hematopoietic origin of the profiled cells (**Supplemental Figure 2D**). Differential gene expression analysis across all splenocyte clusters identified transcriptional changes, including upregulation of Tgfbr1, Fyb and Ifi208 (Pydc3), alongside decreased expression of Ccnb1ip1, Prkch, Abtb2 and Tox (**Supplemental Figure 2E**). Visualization of Tgfbr1 expression across all annotated splenic immune populations demonstrated reproducible elevation in multiple cell subsets from AIF1-cKO mice across biological replicates (**Supplemental Figure 2F**). Cell-specific analysis revealed greatest changes in differential gene expression within DC, memory T cell, neutrophils and pre/pro-B cell populations (**Supplemental Figure 2G**), with negligible changes in mature B cells and naive CD8^+^ or CD4^+^ T cells subsets. Together, these findings indicate that AIF1 deficiency is associated with selective transcriptional remodeling across multiple immune populations, with the greatest changes observed in DC and antigen-experienced or developing immune cell compartments.

### Dendritic cell alterations in the absence of AIF1 are characterized by transcriptional remodeling and reduced cDC1 and pDC frequencies

Given the prominent transcriptional remodeling observed in DC populations and the impaired antigen-specific T cell responses during infection, we conducted differential gene expression analysis within the DC cluster at steady state. scRNA-seq analysis of total splenic DC revealed increased Tgfbr1 expression and reduced Aif1 expression in AIF1-cKO mice compared with controls (**Figure 3A-B**). To resolve changes within DC subpopulations, cDC1, cDC2a, cDC2b, inflammatory monocyte-derived DC (MoDC) and plasmacytoid DC (pDC) subsets were annotated using canonical markers Irf8, Irf4, Zbtb46, Cd24a, Sirpa, Siglech, Tbx21 (T-bet), Klf4 and Ccr2 (**Figure 3C, Supplemental Figure 3**). Aif1 expression was largely confined to cDC1, with minor expression in MoDC and cDC2a, and minimal-to-no expression in cDC2b or pDC subsets (**Figure 3D**).

**Figure 3.**
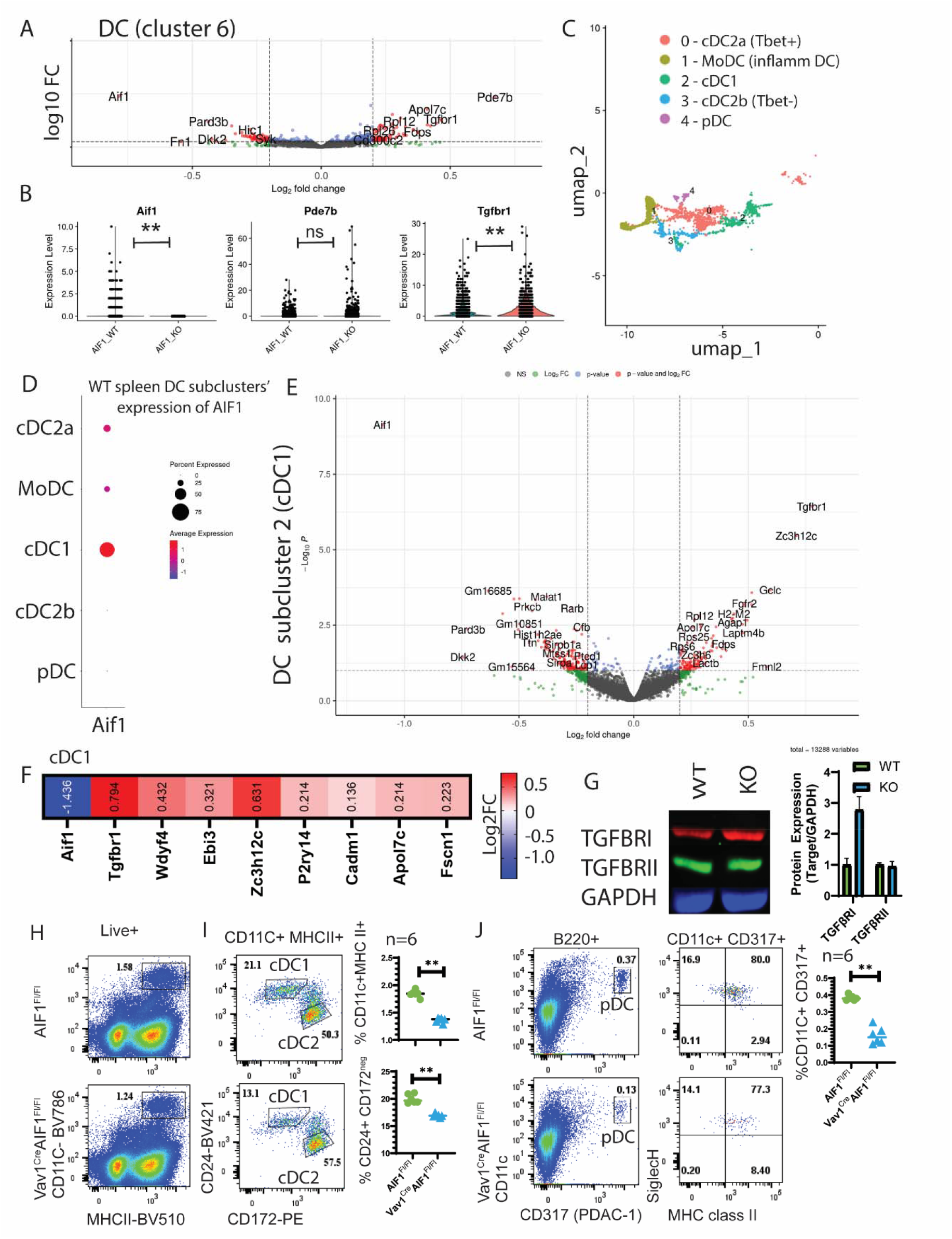
scRNA-seq and flow cytometry analysis of dendritic cells in AIF1-deficient mice. (**A**) UMAP plots showing integrated cell distribution from steady-state spleen scRNA-seq of control (AIF1^fl/fl^) and AIF1-cKO (Vav1^iCre^AIF1^fl/fl^) mice. (**B**) Violin plots showing representative differentially expressed gene Aif1, Pde7b and Tgfbr1. (**C**) Annotation of DC subclusters using canonical markers (Irf8, Irf4, Zbtb46, Cd24a, Sirpa, Siglech, Tbx21, Klf4, Ccr2), identifying cDC1, cDC2a, cDC2b, monocyte-derived DC (MoDC) and plasmacytoid DC (pDC) populations. (**D**) Dot plot showing Aif1 expression across DC subclusters in control mice. (**E-F**) Volcano plot and heat map of differential gene expression analysis of cDC1 cells from AIF1-cKO versus control mice. (**G**) Protein expression in splenic cDC showing for TGFβ receptor I (TGFβRI), TGFβ receptor II (TGFβRII) and GAPDH in AIF1-cKO mice compared with controls. (**H**) Frequency of total CD11c+MHC class II+ DC in spleen from control and AIF1-cKO mice as analyzed by flow cytometry. (**I**) Frequency of CD24+CD172a^neg^ cDC1 subset in spleen. (**J**) Frequency of B220+CD11c+CD317+ pDC in spleen. Data are presented as mean ± SEM; individual points represent biological replicates in bar graphs. Statistical significance is indicated in the corresponding figure panels.

Subset-specific differential expression analysis revealed distinct transcriptional programs across cDC subsets. In cDC1 from AIF1-cKO mice, Tgfbr1 expression was increased, together with altered expression of genes involved in immune regulation and cellular signaling, including Zc3h12c, Ebi3, Apol7c and Cadm1 (**Figure 3E-F**). T-bet^+^ cDC2a cells also exhibited increased Tgfbr1 and Cxcl9 expression, together with reduced expression of genes involved in cell cycle and metabolic regulation, including Ccnb1ip1, Tagln2 and Pfkb3 (**Supplemental Figure 3A-B**). Despite minimal baseline Aif1 expression, T-bet^neg^ cDC2b cells exhibited a similar pattern, with increased Tgfbr1 and reduced expression of transcriptional and signaling regulators, including Junb, Id2, Cdk8 and Lars2 (**Supplemental Figure 3C-D**). In contrast, pDC showed a distinct transcriptional profile, with no significant change in Tgfbr1, but increased expression of Fyb, Pgam2 and Ddx5, alongside decreased Ern1, Maml3, and Prkar2b (**Supplemental Figure 3E-F**).

Consistent with the scRNA-seq findings, western blot analysis of MACS-enriched splenic CD11c^+^ cDC confirmed increased TGFβ receptor I (TGFβRI) protein expression in AIF1-cKO mice compared with controls, whereas TGFβ receptor II (TGFβRII) expression was unchanged (**Figure 3G**).

To further define DC alterations associated with AIF1 deficiency, we assessed cDC and pDC subset frequencies by flow cytometry at steady state. Total CD11c^+^MHC class II^+^ DC were reduced in AIF1-cKO mice, with marked decreases in both cDC1 and pDC populations (**Figure 3H**). Subset analysis further identified reduced frequencies of CD24^+^CD172a^neg^ cDC1 (**Figure 3I**) and B220^+^CD11c^+^CD317^+^ pDC populations (**Figure 3J**). Thus, AIF1 deficiency is associated with both transcriptional remodeling of multiple DC subsets and reduced representation of cDC1 and pDC populations at steady state.

### Loss of AIF1 is associated with altered macrophage transcriptional and functional states

To determine the baseline transcriptomic contributions of AIF1 in resting immune populations, we analyzed splenic macrophages from WT and AIF1-cKO mice by scRNA-seq (**Figure 4A**). Differential expression analysis of the broader macrophage population identified a limited set of transcriptional changes, including increased expression of Tgfbr1, Rps27rt and Rps25, together with decreased expression of Sox5 and Lyn (**Figure 4B-C**). Unsupervised clustering identified an AIF1^high^ macrophage population that was preserved in AIF1-cKO mice despite efficient deletion of Aif1 transcripts, indicating that the transcriptional state defining this macrophage population persists in the absence of AIF1 (**Figure 4D**). DEG analysis within this distinct AIF1high population revealed limited transcriptional differences, with increased Tgfbr1 expression emerging as the principal recurring feature (**Figure 4E-F**). The AIF1^low^ macrophage cluster had demonstrably little AIF1 expression (**Figure 4G**).

**Figure 4.**
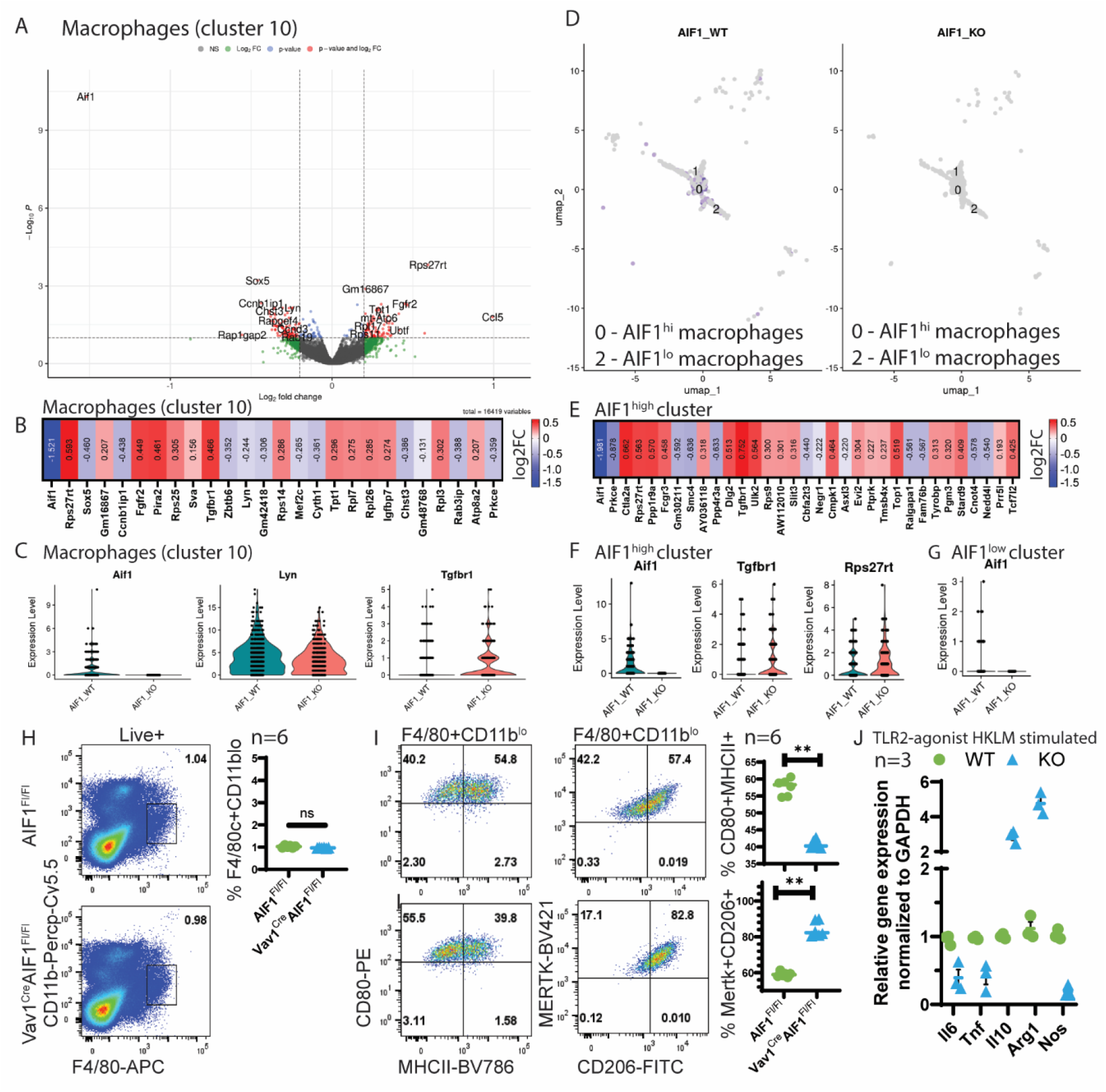
scRNA-seq and flow cytometry analysis of macrophages in AIF1-deficient mice. (**A**) Volcano plot of differentially expressed genes across the total splenic macrophage compartment from control (AIF1^fl/fl^) and AIF1-cKO (Vav1^iCre^AIF1^fl/fl^) mice. (**B-C**) Volcano and violin plots of differentially expressed genes in total macrophages (cluster 10) from scRNA-seq data. (**D**) UMAP feature plot of the distinct macrophage subclusters showing AIF1^high^ (subcluster 0) vs Aif1^low^ (subcluster 2); subcluster 1 was excluded upon identification of B cell contamination. (**E-F**) Volcano and violin plots of differentially expressed genes in AIF1^high^ macrophages. (**G**) Violin plot of Aif1 gene expression in the AIF1^low^ subcluster. (H) Frequency of F4/80+CD11b^lo macrophages. (**I**) Expression of CD80 and MHC class II, and Mertk and CD206 within the gated F4/80^+^CD11b^lo macrophage population. (**J**) qPCR analysis of gene expression in splenic-derived macrophages from control or AIF1-cKO mice stimulated *ex vivo* with the TLR2 agonist (heat-killed Listeria monocytogenes). Data are mean ± SEM. Bar graphs show representative data points across replicates and statistical analyses as indicated in figure panels.

Flow cytometric analysis showed no significant differences in total splenic macrophage frequencies between AIF1-cKO and WT mice. However, phenotypic profiling of the F4/80^+^CD11b^lo^ macrophage compartment at steady state revealed reduced MHC class II expression together with increased co-expression of Mertk and CD206 in AIF1-cKO mice (**Figure 4H-I**), indicating an altered macrophage activation phenotype with increased expression of immunoregulatory-associated markers.

To evaluate whether these steady-state differences translated into altered innate responsiveness, isolated splenic macrophages were stimulated *ex vivo* with the TLR2 agonist HKLM. Following stimulation, AIF1-cKO macrophages exhibited reduced expression of Il6, Tnf, and Nos2, together with increased expression of Il10 and Arg1 (**Figure 4J**). Together, these findings demonstrate that AIF1 deficiency is associated with selective transcriptional remodeling of splenic macrophages, altered expression of activation-associated surface markers and a reduced inflammatory response to TLR2 stimulation.

### Loss of AIF1 in antigen-presenting cells is associated with cell-extrinsic transcriptional remodeling of T cell populations

Given that AIF1 expression was restricted to myeloid APC, scRNA-seq analysis was performed to examine whether AIF1 deficiency was associated with broader transcriptional alterations within naive and memory T cell populations. Although AIF1 expression was absent from T cells, AIF1-cKO mice exhibited increased Tgfbr1 and Fyb expression within memory T cell populations at steady state (**Supplemental Figure 5A-B**). Naïve T cells showed fewer transcriptional changes overall, but exhibited increased expression of Tgfbr1 and S100a10, together with reduced expression of Sell (CD62L) and Pdcd4 (**Supplemental Figures 5C-D**). To determine whether elevated Tgfbr1 expression was reproducibly associated with AIF1 deficiency, individual biological replicates were analyzed independently. Increased Tgfbr1 expression was consistently observed across DC, macrophages and both naive and memory T cell populations from AIF1-cKO mice in each biological replicate, demonstrating that this transcriptional signature was not driven by a single outlier sample (**Supplemental Figure 5E**). These findings indicate that AIF1 deficiency in myeloid APC is associated with transcriptional alterations that extend beyond the AIF1-expressing compartment, affecting both naïve and memory T cell populations.

### AIF1 deficiency enhances TGFβ responsiveness and contributes to impaired myeloid immune function

Given the recurrent increase in Tgfbr1 expression associated with AIF1 deficiency, we examined whether altered TGFβ receptor signaling contributed to the functional defects observed in AIF1-deficient myeloid APC. CRISPR/Cas9-mediated reduction of Tgfbr1 achieved >68% knockdown efficiency in DC, enabling assessment of receptor contribution to functional responses (**Figure 5A**). In DC-T cell co-culture assays, AIF1-cKO DC exhibited reduced capacity to support CD8+ T cell proliferation compared with WT DCs. Tgfbr1 knockdown in AIF1-cKO DCs partially restored CD8+ T cell proliferation, indicating that increased Tgfbr1 expression contributes to, but does not fully account for, the functional impairment observed (**Figure 5B**). To evaluate immune responsiveness in a physiologic inflammatory context, total splenic cells from AIF1-cKO mice were subjected to Tgfbr1 silencing and stimulated with HKLM. 18 h after stimulation, splenic macrophages were isolated by magnetic bead enrichment and analyzed by qPCR for inflammatory cytokine expression. AIF1-cKO macrophages from the Tgfbr1-silenced cohort exhibited partial restoration of Il6, Tnf and Il10 expression (**Figure 5C**).

**Figure 5.**
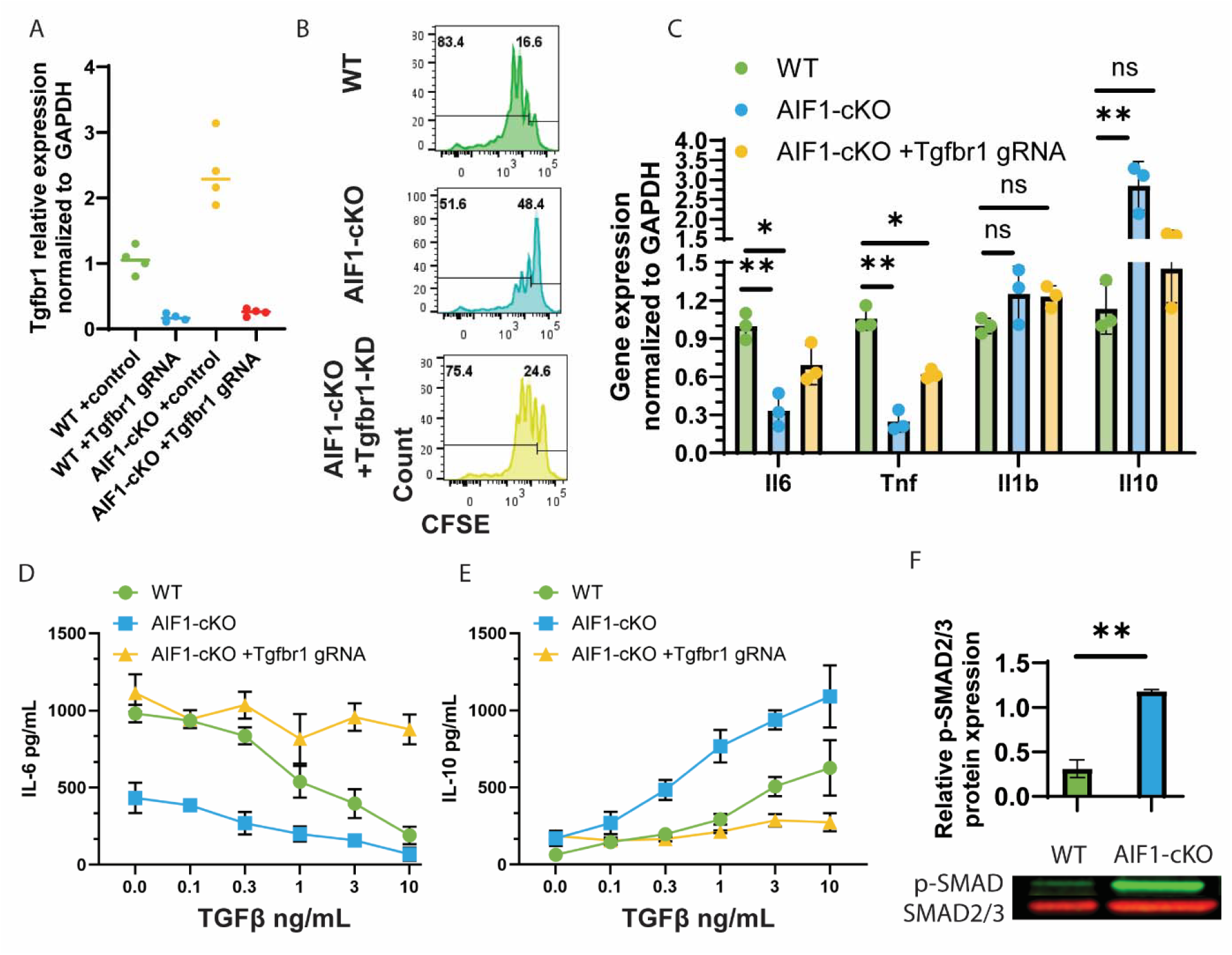
AIF1 deficiency is associated with enhanced TGFβ responsiveness and enhanced SMAD signaling activation *in vitro*. (**A**) Relative mRNA expression (qPCR) of Tgfbr1 in DC following CRISPR/Cas9-mediated silencing to confirm knockdown efficiency. (**B**) Proliferation of CFSE-labeled CD8+ T cells in co-culture with WT or AIF1-cKO BMDC following Tgfbr1 silencing or control treatment and stimulation with LM-OVA. (**C**) Relative mRNA expression (qPCR) of Il6, Tnf, Il1b and Il10 upon HKLM treatment of macrophages following CRISPR/Cas9 Tgfbr1 silencing. (**D-E**) Dose-response curves of measuring IL-6 and IL-10 cytokine levels in control and AIF1-cKO pre-treated with increasing concentrations of recombinant TGFβ1 prior to HKLM stimulation. (**F**) Western blot analysis of phosphorylated SMAD2/3 and total SMAD2/3 levels in control and AIF1-cKO myeloid cells following stimulation with recombinant TGFβ1. Data are mean ± SEM; bar graphs show representative data points across replicates and statistical analyses as indicated in the figure panels.

Next, the study evaluated whether increased Tgfbr1 expression was associated with enhanced responsiveness to TGFβ1. Total splenocytes from AIF1-cKO demonstrated reduced IL-6 production and increased IL-10 induction in response to lower levels of administered TGFβ1 compared with WT controls **(Figure 5D-E)**, indicating enhanced TGFβ responsiveness. To evaluate downstream pathway engagement, phosphorylation of SMAD2/3, a key mediator of canonical TGFβ signaling, was assessed following low-dose TGFβ administration *in vitro*. AIF1-deficient cells exhibited increased phospho-SMAD2/3 relative to WT controls (**Figure 5F**). Together, these data identify enhanced TGFβ responsiveness as a functional consequence of AIF1 deficiency that contributes to, but does not fully explain, impaired immune function.

## Discussion

Collectively, our findings support a model in which AIF1 functions as a hematopoietic immune-readiness factor that helps establish the cellular conditions necessary for effective innate and adaptive immune responses during infection. Loss of AIF1 impaired control of *Listeria monocytogenes*, reduced pathogen-specific T cell expansion and was associated with a shift in immune activation states toward reduced inflammatory responsiveness and enhanced regulatory features. Rather than identifying a single defective pathway, our results demonstrate that AIF1 influences APC composition, cellular activation states and inflammatory responsiveness across the hematopoietic compartment, thereby shaping the capacity of the immune system to mount effective responses to challenge.

Among myeloid APC populations, cDC1 emerged as one of the subsets most prominently affected by AIF1 deficiency. This finding is consistent with the preferential expression of AIF1 within cDC1 and the central role of this subset in cross-presentation and CD8^+^ T cell priming. Notably, both cDC1 and pDC frequencies were reduced despite Aif1 expression being largely restricted to cDC1 at steady state. Given the close developmental relationship between these lineages and their shared dependence on IRF8-directed differentiation programs, this observation raises the possibility that AIF1 influences a shared developmental or homeostatic process within the IRF8-dependent DC axis, although the specific basis for the reduction in these populations remains unresolved. In contrast, macrophage frequencies were largely preserved despite selective transcriptional and phenotypic remodeling, suggesting that AIF1 differentially influences lineage maintenance and functional programming across myeloid populations. Macrophages remain present but exhibit attenuated inflammatory effector function, including reduced production of pro-inflammatory cytokines and a shift toward immunoregulatory output following innate stimulation, suggesting altered macrophage inflammatory responsiveness in the absence of AIF1. Together, these findings identify distinct alterations in dendritic cell composition and macrophage functional state that are likely to contribute to the impaired pathogen-specific T cell responses observed during infection.

A notable finding was the presence of reproducible transcriptional alterations within naïve and memory T cell populations, despite AIF1 expression being restricted to myeloid APC. This observation suggests that the effects of AIF1 deficiency extend beyond cell-intrinsic mechanisms and instead reflect remodeling of the broader immune environment. Whether these effects arise during lymphocyte development within primary lymphoid organs or through persistent conditioning by AIF1-deficient APC within secondary lymphoid tissues remains unresolved. However, the detection of transcriptional alterations in both naïve and memory T cell compartments at steady state suggests that immune remodeling is established before experimental infection and may reflect persistent consequences of altered myeloid cell function. Importantly, these changes occurred in the absence of major shifts in T cell subset frequencies, indicating that altered cellular conditioning and activation-associated transcriptional states may precede overt changes in immune cell composition.

One of the most consistent transcriptional signatures identified across affected immune populations was increased expression of Tgfbr1. We do not interpret this finding to indicate that TGFβ signaling is the primary mechanism through which AIF1 regulates immunity. Rather, elevated Tgfbr1 expression appears to represent one feature of a broader shift in immune conditioning associated with AIF1 deficiency. The recurrence of this signature in both AIF1-expressing and -non-expressing populations is consistent with a broader cell-extrinsic component to the immune remodeling associated with AIF1 deficiency. More broadly, the collective phenotypes observed across DC, macrophages and T cells support the concept that loss of AIF1 alters baseline immune activation states and responsiveness, favoring a less inflammatory and more restrained response state in selected immune populations. Consistent with this interpretation, *ex vivo* studies demonstrated enhanced responsiveness to TGFβ stimulation, increased SMAD2/3 activation and partial functional restoration following Tgfbr1 silencing. These findings establish biological relevance for the pathway while supporting the conclusion that altered TGFβ responsiveness is one measurable consequence of AIF1 deficiency within a larger program of immune remodeling.

Previous studies have implicated AIF1 in cytoskeletal organization, migration, phagocytosis, and inflammatory signaling through pathways including PKC, MAPK, and NFκB^12, 13, 15, 17, 38-41 11^. The present study extends these observations by demonstrating that disruption of AIF1 has consequences for coordinated immune responses at the organismal level during infection. Importantly, these findings provide functional context for prior human genetic and expression studies linking AIF1-associated loci to autoimmune and inflammatory disease. Rather than acting solely as a marker of myeloid cells, AIF1 appears to influence the functional state of myeloid APCs and the adaptive immune responses that emerge from those states.

Collectively, the data support a model in which AIF1 acts as an integrator of myeloid immune preparedness, helping establish the activation states and response capacity of APCs required for effective inflammatory responses, antigen presentation and adaptive immune priming. In this model, AIF1 sustains the inflammatory competence of APCs and enables effective adaptive immune priming during infection, while loss of AIF1 during hematopoietic development is associated with altered immune activation states and reduced functional host defense.

## MATERIALS AND METHODS

### Sex as a biological variable

Both male and female mice were used in all *in vivo* and *ex vivo* studies; similar findings are reported for both sexes.

### Animals

Mouse models included C57BL/6 (WT, Jackson), AIF1^fl/fl^ (B6.129P-Aif1tm17mwl), Vav1iCre (B6.Cg-Tg(Vav1iCre)A2Kio/J), AIF1^fl/fl^ mice were generated in-house on a C57BL/6 background by CRISPR/Cas9-directed insertion of loxP sites flanking exons 3 through 7 of the Aif1 gene. Mice were subsequently crossed to Vav1iCre animals to achieve hematopoietic-specific deletion. Genotyping for floxed Aif1 and Vav1iCre alleles was performed by standard PCR using genomic DNA isolated from tail biopsies. Mice were bred and maintained under specific pathogen-free conditions at the University of Minnesota and monitored by Research Animal Resources (RAR). Both male and female mice, 6-18 weeks of age, were used for experiments. All procedures were approved by the Institutional Animal Care and Use Committee and conducted in accordance with institutional and NIH guidelines.

### Listeria monocytogenes infection and bacterial burden

For infectious challenge experiments, mice were infected intravenously with Listeria monocytogenes expressing ovalbumin (LM-OVA, strain EGY2961) at the indicated doses (1×10^3^–5×10^3^ CFU). At specified time points post-infection, spleens and livers were harvested, homogenized in sterile PBS, serially diluted, and plated on brain-heart infusion agar plates. Colony-forming units (CFU) were enumerated after overnight to 48 h incubation at 37°C. Serum was collected by cardiac puncture for cytokine quantification.

### Antigen-specific T cell responses

Listeria-specific CD8+ and CD4+ T cell responses were quantified using peptide:MHC (pMHC) tetramers to directly enumerate pathogen-specific T cells *in vivo*. H-2Kb–SIINFEKL tetramers were used to detect OVA_257–264_-specific CD8+ T cells and I-Ab tetramers loaded with LLO- or OVA-derived peptides were used to identify LM-specific CD4+ T cells. Single-cell suspensions from spleen were prepared at day 3-5 post-infection, stained with pMHC-APC tetramers followed by enrichment of APC-positive cells using magnetic bead sorting. The cells were stained with surface markers (i.e. CD3, CD4, CD8, CD44, CD62L, CD122, PD⍰1) and acquisition on a BD FACSLyric and analysis in FlowJo (BD). Internal controls included CD4+ T cells stained with MHC class I tetramers and CD8+ T cells stained with MHC class II tetramers to verify tetramer specificity and background binding.

### CRISPR/Cas9-mediated silencing of Tgfbr1

*In vitro* silencing of Tgfbr1 was performed using RNP (ribonucleoprotein) complexes. Briefly, 1.5 x 10^6^ bone marrow-derived DC or macrophages were resuspended in transfection solution (MEM medium with no antibiotics, FBS) containing Cas9 nuclease complexed with synthetic sgRNA targeting the murine Tgfbr1 locus (Target sequence: GCAGCTCCTCATCGTGTTGG, CATACAAACGGCCTATCTCG) at 1:1 ratio.

Electroporation was performed using the ECM squarewave electroporator (BTX/Harvard Apparatus) with a single low-voltage pulse at 230 V using parameters optimized for primary myeloid cells. Knockdown efficiency was validated 24-48 hours post-electroporation via qPCR for Tgfbr1 mRNA levels relative to scramble gRNA and vehicle-only controls.

### DC-T cell co-culture and proliferation assay

DC-T cell proliferation assays were performed using either Flt3L-generated bone marrow-derived DC (BMDC) or splenic-derived DC isolated from WT and AIF1-cKO mice by magnetic bead enrichment (Miltenyi Biotec). In both systems, DC were subjected to CRISPR/Cas9-mediated Tgfbr1 silencing prior to antigen loading. DC were pulsed with heat-killed Listeria monocytogenes expressing ovalbumin (HKLM-OVA) for 4 h and washed extensively. CFSE-labeled CD8+ T cells isolated from OT-I transgenic mice were then co-cultured with DC for 72-96 h at a 1:10 (DC:T) ratio, as indicated. T cell proliferation was assessed by CFSE dilution using flow cytometry.

### *Ex vivo* splenocyte stimulation and macrophage isolation

Total splenocytes from WT and AIF1-cKO mice were cultured in complete RPMI and stimulated with heat-killed Listeria monocytogenes (HKLM). In selected experiments, splenocytes were subjected to CRISPR/Cas9-mediated Tgfbr1 silencing prior to stimulation. At defined early and late post-stimulation time points, macrophages were isolated by magnetic bead enrichment using anti-F4/80 microbeads (Miltenyi Biotec). Purified macrophages were processed for RNA isolation and quantitative PCR analysis of inflammatory and regulatory gene expression, including Il6, Tnf, Il10, Nos2, and Arg1.

### Phospho-SMAD2/3 analysis

Single-cell suspensions were stimulated with recombinant murine TGFβ1 at the indicated concentration and time points prior to fixation with paraformaldehyde. Cells were permeabilized using methanol-based intracellular staining protocols and stained with phospho-SMAD2/3-specific antibodies. Data were acquired by flow cytometry and analyzed using FlowJo software.

### Antibodies and Bioreagents

The following antibodies (target [clone]) were used for flow cytometry and/or cell sorting: Dendritic cells and myeloid cells: CD11c (N418), MHC class II (M5/114.15.2), CD24 (M1/69), CD172a/SIRPα (P84), CD11b (M1/70), F4/80 (BM8), CD64 (X54-5/7.1), CD317/PDCA-1 (eBio927), B220 (RA3-6B2), SiglecH (eBio440C), Ly6G (1A8), Ly6C (HK1.4), CD80 (16-10A1), CD86 (GL-1), CD40 (1C10), CD206 (C068C2). T cells: CD3 (17A2), CD4 (RM4-5), CD8α (53-6.7), CD25 (PC61), CD44 (IM7), CD62L (MEL-14), CD69 (H1.2F3), PD-1 (29F.1A12), CD122/IL-2Rβ (TM-β1). Intracellular/transcription factors: IRF8 (V3GYWCH), IRF4 (3E4), Foxp3 (FJK-16s).Fluorochrome-conjugated antibodies were purchased from BioLegend, BD Biosciences, eBioscience, and Thermo Fisher. AIF1/Iba1 antibodies were purchased from Abcam. Recombinant cytokines (GM-CSF, Flt3L, M-CSF) were obtained from PeproTech/Thermo Fisher Scientific.

### Single-Cell RNA Sequencing

Spleens from age- and sex-matched WT and AIF1-cKO mice at steady state were dissociated into single-cell suspensions using a GentleMACS tissue dissociator (Miltenyi Biotec), followed by ACK lysis for red blood cell removal. Cell suspensions were filtered through a 70 μm strainer. Approximately 50,000 live cells per sample were sorted and loaded onto the 10x Genomics Chromium platform. cDNA libraries were prepared using 5’ chemistry (10x Genomics) according to manufacturer instructions and sequenced on an Illumina NovaSeq.

Raw reads were processed and aligned to the mm10 mouse reference genome using CellRanger (10x Genomics). Downstream analyses were performed in R using Seurat v4. Low-quality cells and doublets were removed based on UMI counts, gene number and mitochondrial gene content. Data were normalized, scaled and integrated across samples. Dimensionality reduction was performed by principal component analysis and visualized using UMAP. Differential gene expression between genotypes within defined clusters was assessed using the Wilcoxon rank-sum test. Gene set enrichment analyses were performed using MSigDB hallmark gene sets. Cell type annotation was based on canonical marker expression and published reference datasets.

### Flow cytometry and cell sorting

Single-cell suspensions from spleen or lymph node were prepared in PBS containing 0.5% BSA and 2 mM EDTA (MACS buffer). Red blood cells were lysed with ACK buffer where indicated, and cells were stained with the indicated panels of fluorochrome-conjugated antibodies on ice and washed in FACS buffer. For intracellular staining (e.g., IRF8, IRF4, transcription factors), cells were fixed and permeabilized using commercial fixation/permeabilization solutions or 3% paraformaldehyde followed by 0.15% saponin. Data acquisition was performed on BD FACSLyric or BD FACSVerse instruments.

### Cytokine Analysis (Luminex and ELISA)

Serum cytokines were quantified using a multiplex Luminex assay (Thermo Fisher Scientific, Biotechne) for IFNγ, IL-1β, IL-4, IL-6, IL-10, IL-12p70, TGFβ1, and TNFα according to manufacturer protocols. Briefly, samples were thawed on ice, diluted, incubated with capture beads, followed by biotinylated detection antibodies. Beads were acquired on a Magpix analyzer and cytokine concentrations were calculated from standard curves.

### RNA isolation and quantitative PCR

For validation assays, RNA was extracted with TRIzol reagent (Thermo Fisher), reverse-transcribed into cDNA and analyzed by quantitative PCR (qPCR). Reactions were run using TaqMan master mix and probe sets for candidate genes. Housekeeping controls included GAPDH and β-actin. Data were normalized using the ΔΔCt method.

### Western blotting

Protein lysates were generated from cells or tissues in lysis buffer containing protease and phosphatase inhibitors and separated on 10% SDS-PAGE gels. Proteins were transferred to nitrocellulose membranes and probed with respective antibodies. Primary antibodies were detected using fluorescent-labeled secondary antibodies and visualized on a LI-COR Odyssey imaging system (Licor Biosciences).

### Statistical Analyses

Statistical analyses were performed using GraphPad Prism. For comparisons between two groups, unpaired two-tailed Student’s t-tests were used. For comparisons involving more than two groups, one-way ANOVA or two-way ANOVA with appropriate post hoc tests were applied as indicated. Data are presented as mean ± SEM unless otherwise noted. P values < 0.05 were considered statistically significant.

## Acknowledgements

We thank the University of Minnesota Genomics Center for single-cell RNA sequencing library preparation and Illumina NovaSeq sequencing services; the University of Minnesota Supercomputing Institute for high-performance computing resources and bioinformatics support enabling scRNA-seq data processing and Seurat analysis; and the University of Minnesota Flow Cytometry Core for expert assistance with multicolor flow cytometry, cell sorting, and tetramer staining. We are also grateful to Dr. Mark Jenkins (Center for Immunology, University of Minnesota) for providing Listeria monocytogenes expressing ovalbumin and for supplying peptide-MHC tetramers, and to the NIH Tetramer Core Facility for additional peptide-MHC tetramers used in these studies. Core services were supported by the University of Minnesota’s institutional commitment to advancing biomedical research through state-of-the-art shared instrumentation and expertise. Work herein is supported by NIH NIGMS R35GM145290 and NIH NIAID R21AI168668 both to M. Lipscomb, PI/Project Leader.

## Data and Material Availability

Data supporting the findings of this study are available from the corresponding author upon reasonable request. The single-cell RNA sequencing datasets generated and/or analyzed during the current study will be deposited in the Gene Expression Omnibus (GEO) and will be publicly accessible under accession number GSE314797 within 12 months of publication.

## Author Contributions

Conceptualization, M.W.L., L.R., and R.L.d.S.; methodology, L.R., R.L.d.S., A.R., J.S., and M.M.; investigation, L.R., R.L.d.S., A.R., and J.S.; bioinformatic analysis, L.R., R.L.d.S., and M.M.; formal analysis, L.R., R.L.d.S., and M.M.; data curation, L.R., and R.L.d.S.; supervision, M.W.L.; visualization, L.R., A.R., T.d.M., R.L.d.S.; writing—original draft, M.W.L., L.R., T.d.M., and R.L.d.S.; writing—review and editing, all authors.

