## Supplemental Figures for "Allograft Inflammatory Factor-1 regulates immune activation states and is required for effective pathogen-specific T cell immunity during infection"

**Allograft Inflammatory Factor-1 supports myeloid antigen-presenting cell function and adaptive immune priming in vivo during infection**

#Lais Rekowsky[1], #Ricardo Louzada da Silva[1,2], Ayane Resende[1], Jonathan Seenarine[1], Marissa Macchietto[3], Tatiana de Moura[4], Diana M. Elizondo[5] and Michael W. Lipscomb[1,6]

**Supplemental Figures S1-5**

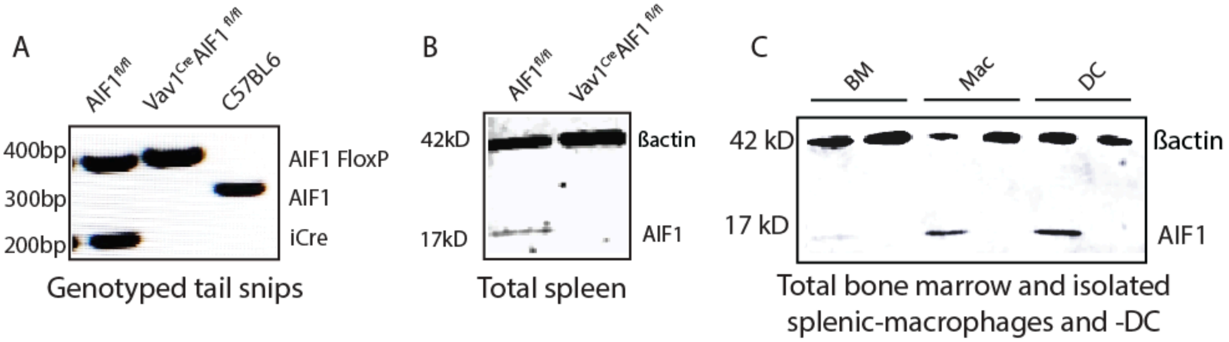


**Supplemental Figure 1. Generation and validation of hematopoietic AIF1 conditional knockout mice.** (**A**) PCR genotyping of floxed AIF1 allele and Vav1-iCre presence in experimental cohorts. (**B-C**) AIF1 protein expression by western blot in splenocytes, bone marrow cells, splenocyte-derived macrophages and dendritic cells from control and AIF1-cKO mice.

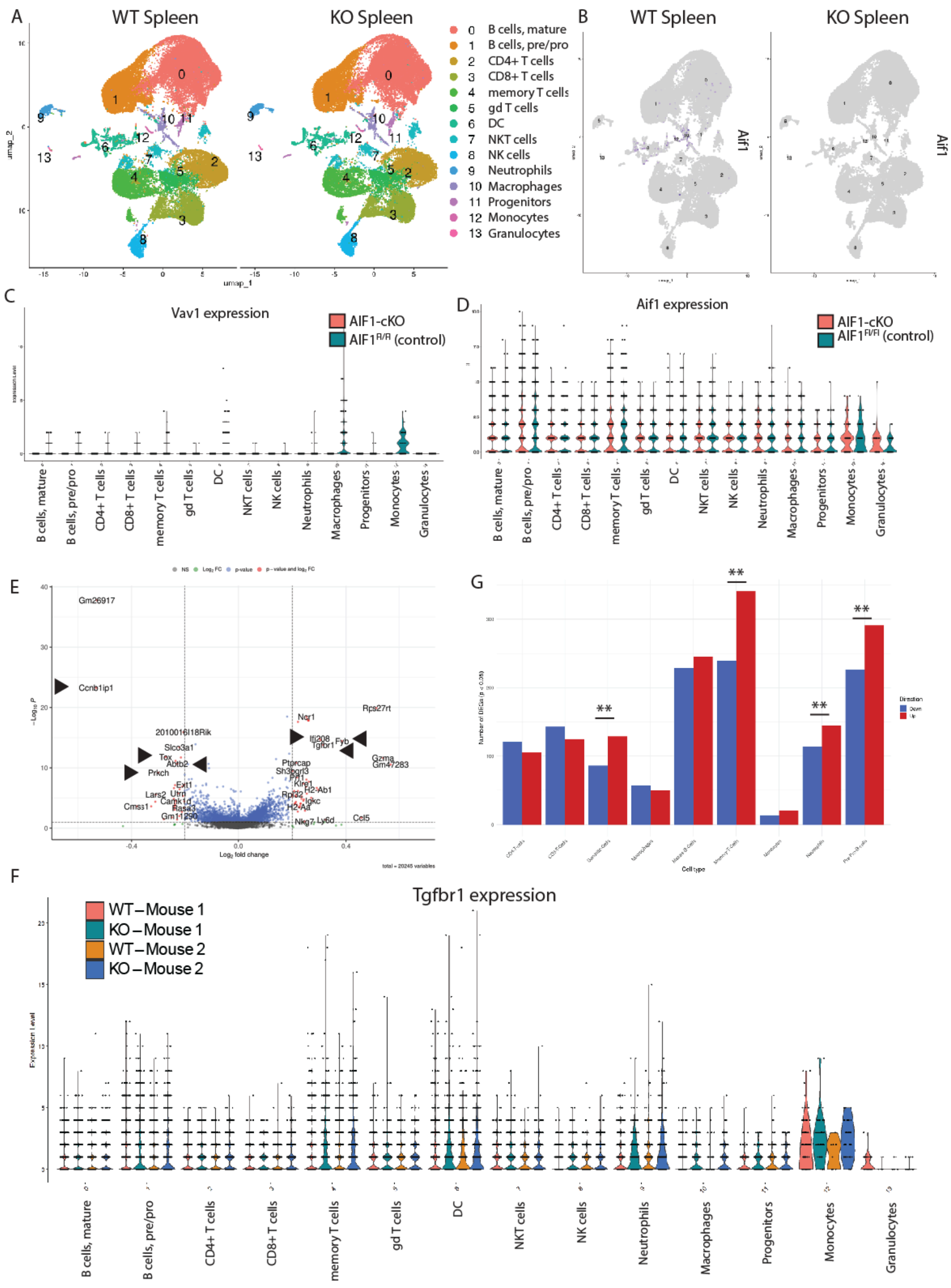


**Supplemental Figure 2. Global scRNA-seq quality control and AIF1 expression patterns at steady state.** (**A**) UMAP plots showing integrated cell distribution from AIF1fl/fl and AIF1-cKO spleen scRNA-seq. (**B**) UMAP feature plot showing Aif1 expression across splenic clusters. (**C-D**) Vav1 and Aif1 expression verification across hematopoietic clusters as shown by violin plots. (**E**) Volcano plot of top differentially expressed genes across all splenocyte clusters. (**F**) Tgfbr1 expression across hematopoietic clusters within independent biological replicates as shown by violin plots. (**G**) Bar graph selecting relative amount of differentially expressed genes across all clusters.

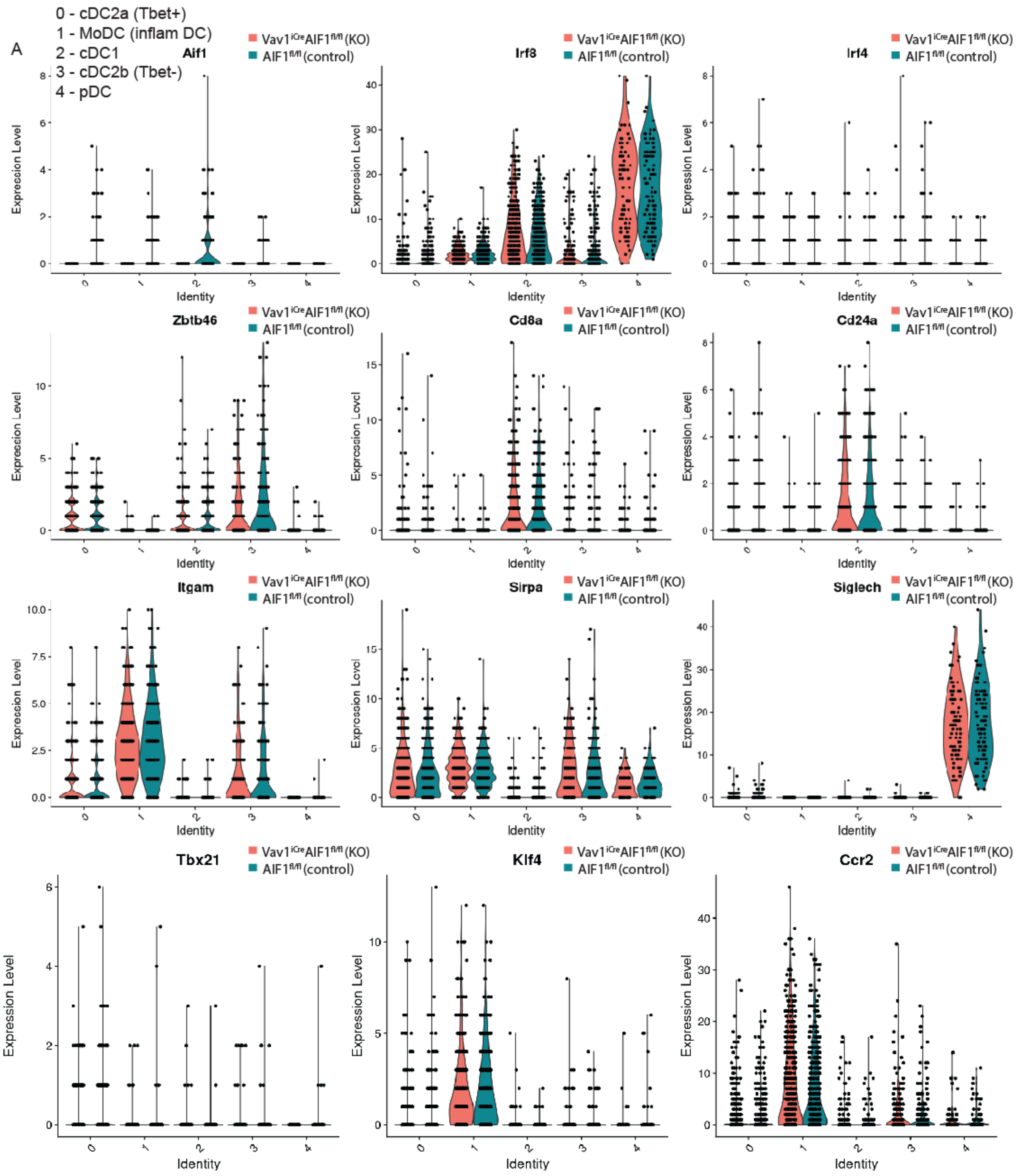


**Supplemental Figure 3. Identification of DC subclusters using canonical markers.** (**A**) UMAP annotation of DC subclusters (cDC1, cDC2a, cDC2b, MoDC, pDC) using canonical markers Irf8, Irf4, Zbtb46, Cd24a, Sirpa, Siglech, Tbx21, Klf4, Ccr2.

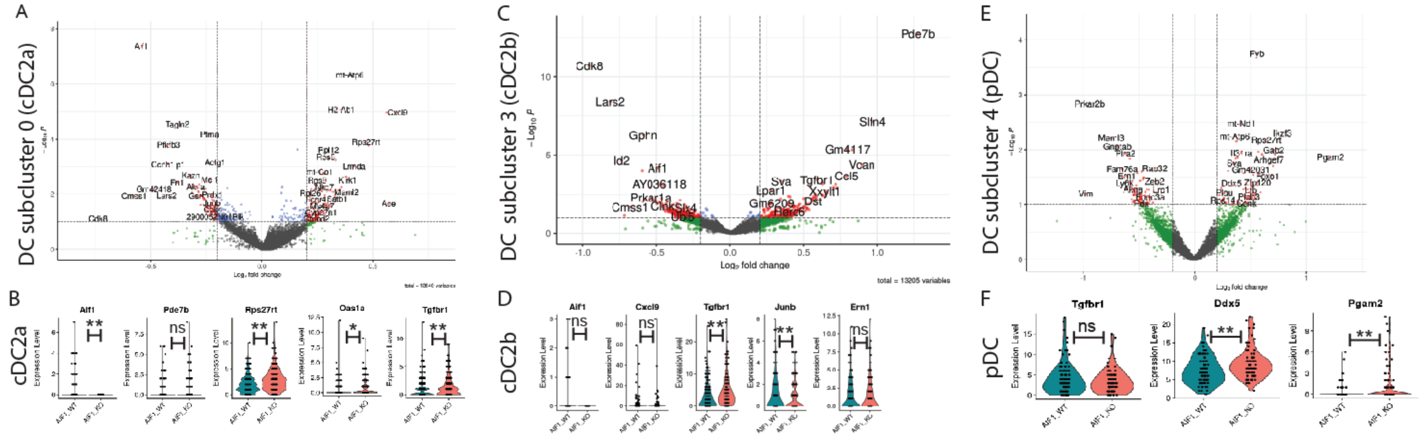


**Supplemental Figure 4. Comprehensive dendritic cell subcluster analysis reveals tolerance-biased programming.** Volcano and violin plots for (**A-B**) cDC2a, (**C-D**) cDC2b and (**F-G**) pDC analysis showing transcriptional change as determined by scRNA-seq.

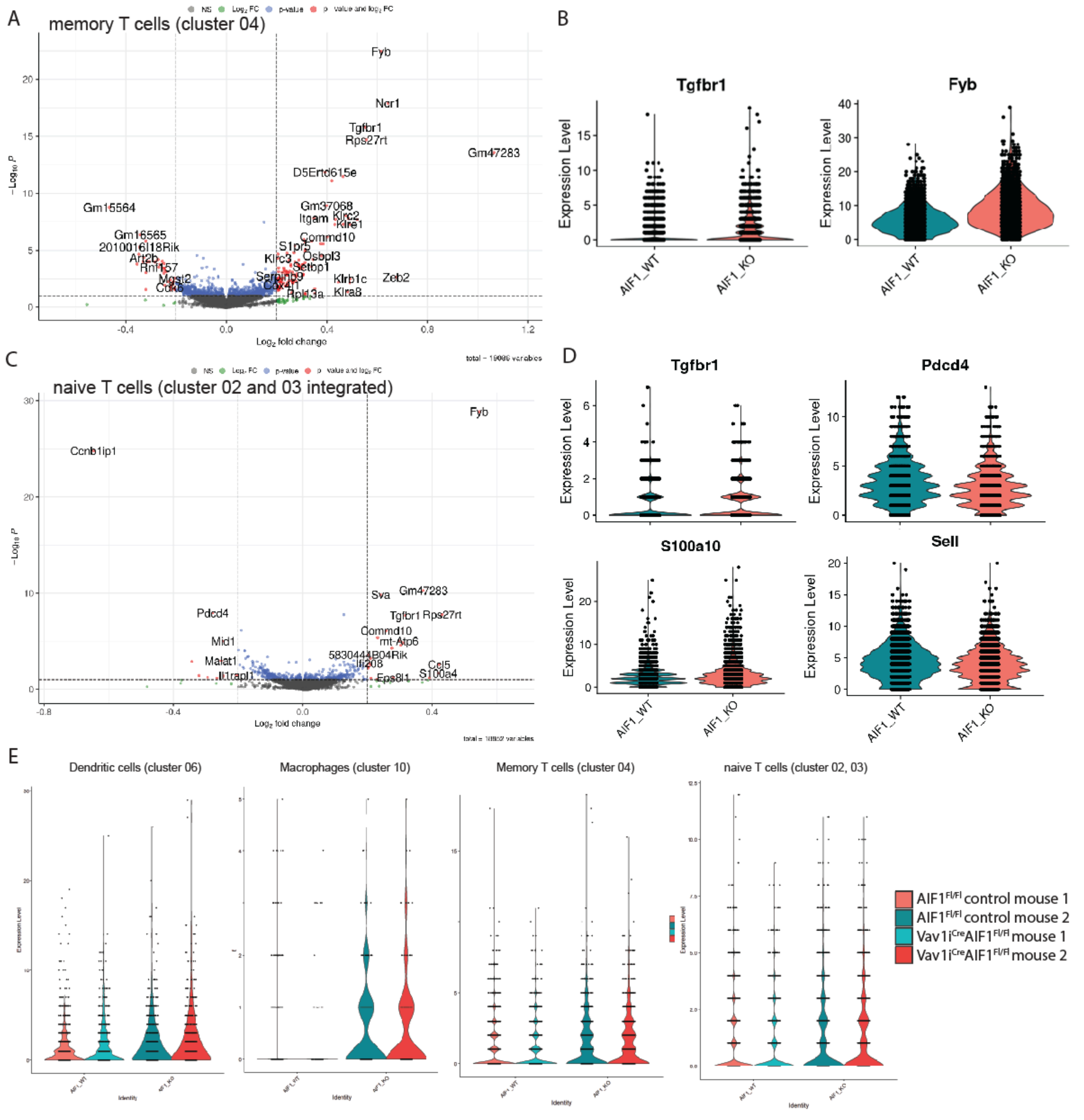


**Supplemental Figure 5. T cell compartment alterations support TGFβ-biased immune programming.** (**A-B**) Volcano and violin plots showing DEG in memory T cell clusters from spleen. (**C-D**) Volcano and violin plots showing differential expression in naive CD8+ and CD4+ T cells cell clusters. (**E**) scRNA-seq expression analysis for levels of Tgfbr1 in DC, macrophages, memory T cells and naive T cells from steady-state spleens of individual biological replicates: AIF1$^{fl/fl}$ (control) mouse 1, AIF1$^{fl/fl}$ (control) mouse 2, AIF1-cKO mouse 1, and AIF1-cKO mouse 2.
